# Movements and habitat use of the invasive blue crab *Callinectes sapidus* in Mediterranean coastal lagoon

**DOI:** 10.1101/2025.08.13.663855

**Authors:** Eric Dominique Henri Durieux, Klervi Le Corre, Guillaume Marchessaux, Dimitri Veyssiere, Sabrina Etourneau, Marie Garrido

## Abstract

The Atlantic blue crab (*Callinectes sapidus*), an invasive and ecologically impactful decapod species, has rapidly proliferated across the Mediterranean Sea, prompting concerns over its effects on native ecosystems and local economies. Despite its growing presence, knowledge of its behavioural ecology in Mediterranean coastal systems remains limited. This study presents the first acoustic telemetry-based assessment of *C. sapidus* within its invasive range, focusing on movement patterns, activity levels, and habitat preferences in Biguglia Lagoon, Corsica (Northwestern Mediterranean). In 2023, 31 adult crabs (20 males, 11 females) were tagged and tracked over four months. Accelerometer data revealed distinct diel activity patterns, with females exhibiting significantly higher walking and swimming behaviours, while males were more sedentary, particularly at night. Salinity preferences varied by sex: females occupied higher-salinity zones, likely linked to reproductive behaviour, while males favoured lower salinity areas. Females also travelled greater distances and had larger, more variable home ranges, with some individuals extending into adjacent marine environments. Although no berried females were tagged, detections at sea suggest early seaward migrations by non-ovigerous and potentially ovigerous females. These findings underscore the species’ behavioural plasticity and sex-specific ecological strategies. The integration of telemetry and environmental data offers critical insights into the spatial ecology of *C. sapidus*, informing targeted management strategies. This research directly supported Corsica’s first Territorial Control Plan for marine invasive species and provides a framework for adaptive management across other Mediterranean regions facing similar ecological challenges.

**Highlights:**

- First telemetry study of blue crab in Mediterranean invasive range
- Females showed higher activity and larger home ranges than males
- Crabs displayed sex-specific salinity and depth preferences
- Some females migrated to sea, suggesting early reproductive movement
- Findings informed Corsica’s first invasive species control plan

## 1. Introduction

The recent report from the Intergovernmental Science-Policy Platform on Biodiversity and Ecosystem Services (IPBES) highlights the alarming global spread of Non-Indigenous Species (NIS) (Roy et al., 2023). Of the 37,000 documented NIS, 3,500 are classified as Invasive Alien Species (IAS), accounting for 60% of global species extinctions. These IAS affect biodiversity by altering ecosystems, competing for resources, and preying on native species (Roy et al., 2023). Climate change intensifies shifts in species distribution, indirectly promoting the spread of IAS, often to the detriment of native species. Tracking both native and invasive species is essential for assessing the impacts of climate change and developing effective conservation measures (Pyšek & Richardson, 2010; Pyšek et al., 2020). Mapping the habitats and behaviours of marine invasive species is crucial for effective management strategies. Traditional data collection methods, limited by observational constraints and small-scale field studies, often fail to capture essential ecological variables like habitat use by new colonizers. To enhance conservation efforts, especially in high-value natural habitats, it is vital to expand the observational scale and incorporate key ecological factors influenced by habitat modulators, such as aggressive NIS (Pyšek et al., 2020). Integrating traditional methods with affordable new technologies can significantly aid conservationists, practitioners, and decision-makers.

The blue crab, *Callinectes sapidus* (Rathbun, 1896), native to the western Atlantic, is known for its remarkable adaptability and colonization rate (Mancinelli et al., 2017; Mancinelli et al., 2021) driven by its high reproductive rate (Hines et al., 1987; Darnell et al., 2009) and broad tolerance to both temperature (Marchessaux et al., 2022) and salinity (Marchessaux et al., 2024a). Since the early 20^th^ century, this species has successfully invaded and thrived in the Mediterranean and Black Sea and along the European coasts of the Atlantic Ocean (Mancinelli et al., 2017; Mancinelli et al., 2021) through successive introduction events via ballast waters (González-Ortegón et al. 2022). Significant outbreaks and northward expansion of blue crab have been recently reported in this invasive range likely triggered by sea water warming (Mancinelli et al., 2021). Notable occurrences include the Northwestern Mediterranean such as Spain (Clavero et al., 2022), Sardinia (Culurgioni et al., 2020), France (Labrune et al., 2019), Corsica (Marchessaux et al., 2024b), as well as the Northern Adriatic (Azzuro et al., 2024), and the Southern Coast of Portugal in the Eastern Atlantic (Vasconcelos et al., 2019). *Calinectes sapidus* is an opportunistic omnivorous and detritic feeder (Laughlin, 1982; Fitz et al. 1992) with aggressive behavior (Hines, 2007). This has led to significant ecological and economic impacts, through predation on a wide range of species, including molluscs, invertebrates, and seagrasses (Prado et al., 2020), and by strongly affecting artisanal fisheries (Clavero et al., 2022; Marchessaux et al., 2023b; Gavioli et al., 2025b).

In the Mediterranean Sea, *C. sapidus* inhabits a variety of productive coastal habitats, such as river mouths, brackish lagoons, and coastal marinas (Mancinelli et al., 2021; Gavioli et al. 2025a), which provide shelter and abundant food resources that support its growth and reproduction (Prado et al., 2020). Thanks to its complex life cycle with multiple larval and juvenile stages, it can thrive in different habitats and across a wide range of salinities (Marchessaux et al., 2024a). Reproduction occurs in low-salinity waters, where females migrate to polyhaline zones to lay their eggs (Aguilar et al., 2005). These eggs develop in seawater, releasing planktonic larvae that eventually return to less saline areas, which serve as nurseries for juveniles (Epifanio, 2019). The blue crab exhibits particularly high larval dispersal and connectivity, which significantly enhances its invasive potential in the Mediterranean (Marchessaux et al., 2023a; Barrier et al., 2025).

Along the French Mediterranean coasts*, C. sapidus* was first recorded in the Berre Lagoon in 1962, and has since gradually extended its range over coastal lagoons of the Gulf of Lions and Corsica especially (Labrune et al. 2019). Between 2014 and 2017, *C. sapidus* expanded along Corsica’s eastern coast (Garrido & Noël., 2018), with a sharp population increase from 2021 to 2024, especially in the Palu and Biguglia lagoons. The first gravid females were reported in Palu in 2020, confirming local reproduction (Unpublished data). These lagoons, both heavily impacted and actively fished, are now closely monitored through ongoing scientific studies (Marchessaux et al., 2023b; 2024a,b; Paganelli et al., 2025), with particular attention from resource managers.

Understanding the movement patterns of specific marine species is crucial for accurately assessing their spatio-temporal dynamics, which is essential for effective management, particularly in the contexts of conservation and fisheries (Hays, 2016). Decapods are among the most successful and influential invaders of freshwater and marine ecosystems, exerting measurable influences on biological communities (Snyder & Evans, 2006, Twardochleb et al. 2013, Matheson et al. 2016). Addressing these challenges requires information on how decapods interact with other ecosystem components across space and time with study of their movements (Florko et al. 2021). Acoustic telemetry is a methodology particularly suited for investigating in situ movements, activity rhythm, home range, territoriality and migration (Arnold & Dewar, 2001; Hussey et al., 2015; Crossin et al., 2017). Several studies in the native range of *C. sapidus* have employed acoustic telemetry to investigate its behavior, particularly focusing on movement patterns, foraging strategies, habitat use, and swimming capabilities (Wolcott & Hines 1990, Hines et al. 1995, Clark et al 1999, Carr et al. 2004, Darnell et al. 2012). Spawning migrations of females from estuarine waters to marine open waters are especially well documented with regards for instance to the influence of tidal currents (Carr et al 2004, Darnell et al. 2012). To date, to our knowledge, no studies have investigated the behavior of *C. sapidus* using acoustic telemetry within its invasive range in the Mediterranean. Given the species’ complex behavioural patterns observed in its native range, studying its movements and habitat use in the Mediterranean lagoon–sea, systems markedly different from its native environments, is particularly relevant.

This study aimed to examine the behaviour of the invasive blue crab in a representative Mediterranean lagoon system, the Biguglia Lagoon in Corsica. Specifically, it focused on activity patterns, movement (distance travelled), and habitat use (home ranges), while also assessing sex-based differences and evaluating the influence of environmental factors, such as salinity, on habitat use.

## 2. Materials and Methods

### 2.1. Study Area

The Biguglia lagoon, located south of Bastia (42°36’N, 09°28’E), is Corsica’s largest coastal lagoon, covering 14.5 km² with a watershed of 180 km² (Figure 1). This brackish lagoon is characterized by shallow waters, with a maximum depth of 1.8 m, and a total volume of 18 Mm³ (Dufresne et al., 2017). Freshwater input is primarily provided by the Bevinco River in the northwest, while the artificial Fossone channel in the south also channels water from the Golo River, depending on river flow, wind, and currents (Figure 1; Erostate et al., 2022). The lagoon connects to the sea through a 1.5 km-long natural sea channel in the north, also known as the grau, which opens intermittently, and through the Fossone channel in the south. The system experiences wide fluctuations in temperature (3°C–28°C) and salinity (4–37 PSU), reflecting its dynamic balance between marine and freshwater influences (Garrido et al., 2016; Ligorini et al., 2022). Recognized for its high biological and ecological value, the Biguglia Lagoon has been designated a RAMSAR wetland since 1991 and a Natural Reserve since 1994. It provides key ecosystem services, including support for small-scale fisheries (Marchessaux et al., 2024b).

**Figure 1.**
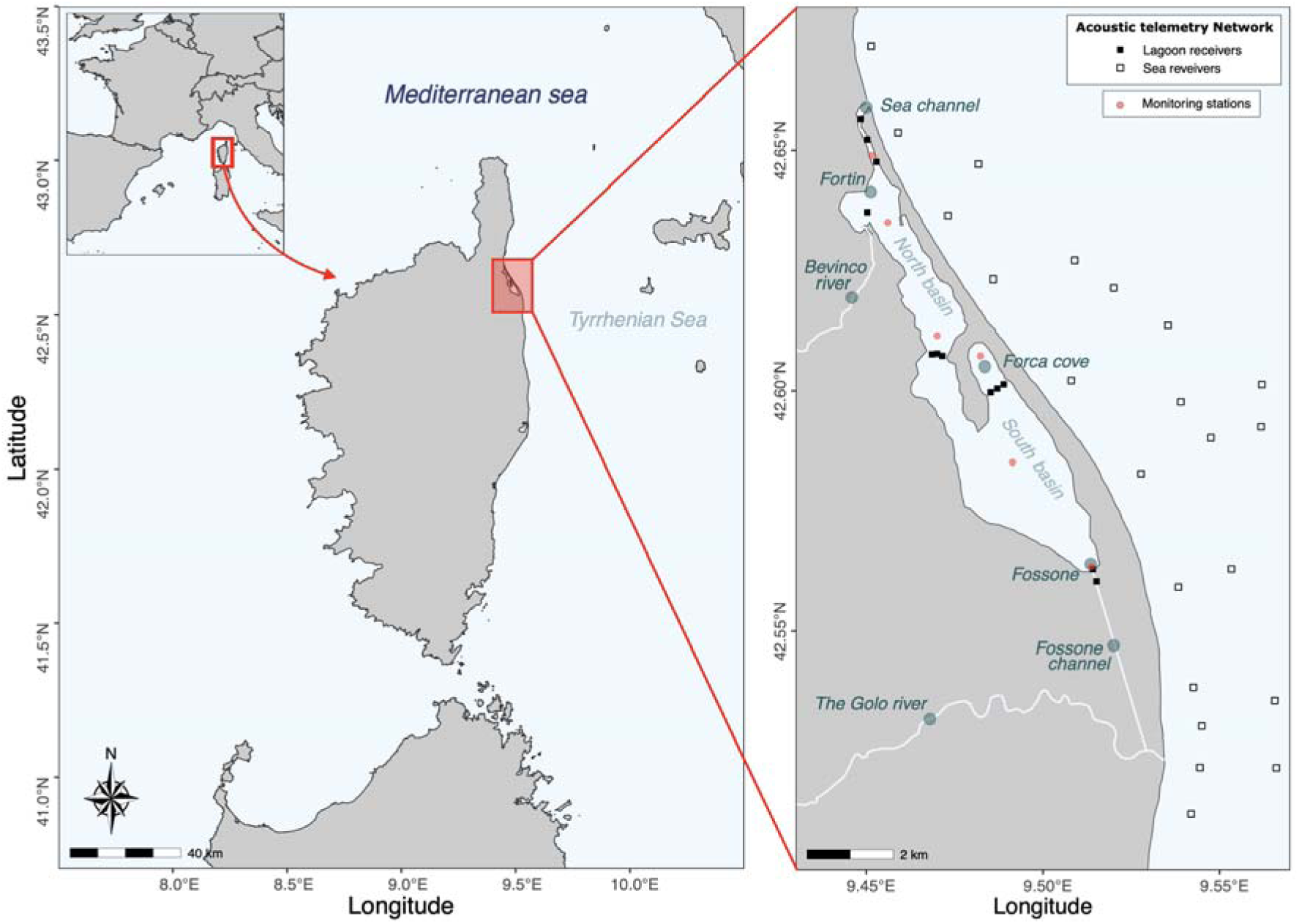
(A) Geographic location of the Biguglia Lagoon within the Northwestern Mediterranean basin. (B) Acoustic telemetry network within the lagoon and adjacent marine area, showing the positions of acoustic receivers - monitoring stations for environmental parameters within the lagoon are also indicated.

### 2.2. Sampling strategy and acoustic telemetry

Three successive catch and tagging campaigns were conducted in 2023 with the help of local fisher using fyke nets. Sampling was distributed across the summer months to fit with spawning season and the overall activity season. It was spread over 3 consecutive months to also to minimize data loss due to individuals moving outside the receiver network, being captured and not retrieved, or experiencing tag detachment, particularly during moulting events. A total of 10, 10, and 11 adult crabs were captured, tagged, and released on the same day in June, July, and August 2023, respectively (Table 1). All individuals were mature - 20 males and 11 females - and none exhibited signs of egg-bearing (Table 1). The crabs were fitted with acoustic transmitters (Supplementary Fig S1.), specifically the Thelma Biotel model ADT-MP9L (Diameter: 9mm; Length 33.7mm; Weight air 5.7 g; Weight water 3.5 g; Power Output 146 dB). The acoustic transmitters emitted at a frequency of 69 kHz (Protocol S64K) at intervals of 30 to 90 seconds (60 sec average). They transmitted depth (m), activity (m.s-2; 3-axis accelerometer with measurements over 1/3 of the average transmit interval) and temperature (°C) (not used in this study) to the receivers on a rolling basis.

**Table 1.**
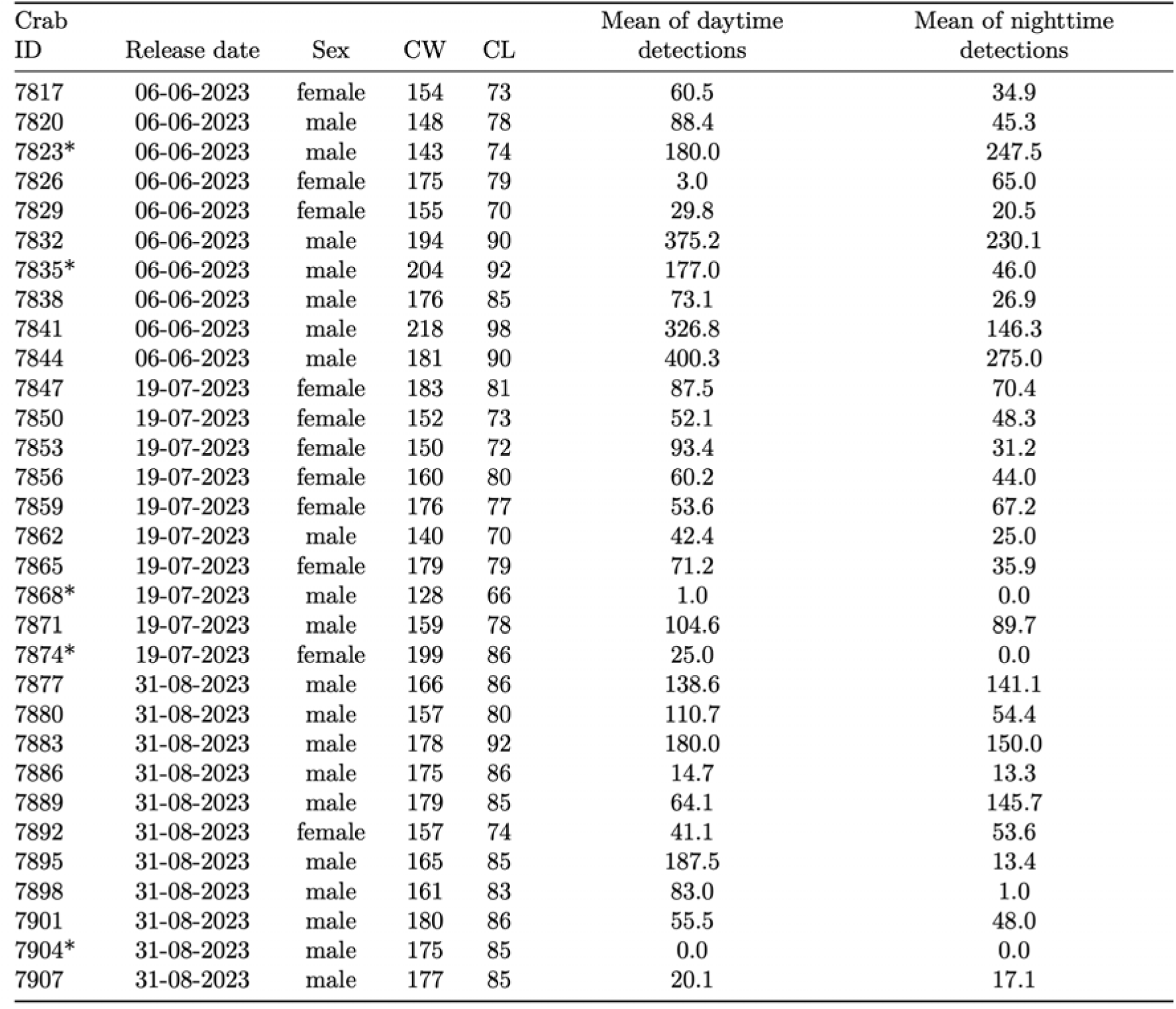
Summary of tagging and detection data for each blue crab (*Callinectes sapidus*) included in the 2023 acoustic telemetry survey in the Biguglia Lagoon (females: n=10; males=20). Note: Individuals marked with an asterisk (*) were excluded from the analysis due to one or more of the following reasons: absence of detections, single detection events, activity recorded only on the day of release, or consistently low activity at a fixed depth.

| Crab ID | Release date | Sex | CW | CL | Mean of daytime detections | Mean of nighttime detections |
| --- | --- | --- | --- | --- | --- | --- |
| 7817 | 06-06-2023 | female | 154 | 73 | 60.5 | 34.9 |
| 7820 | 06-06-2023 | male | 148 | 78 | 88.4 | 45.3 |
| 7823* | 06-06-2023 | male | 143 | 74 | 180.0 | 247.5 |
| 7826 | 06-06-2023 | female | 175 | 79 | 3.0 | 65.0 |
| 7829 | 06-06-2023 | female | 155 | 70 | 29.8 | 20.5 |
| 7832 | 06-06-2023 | male | 194 | 90 | 375.2 | 230.1 |
| 7835* | 06-06-2023 | male | 204 | 92 | 177.0 | 46.0 |
| 7838 | 06-06-2023 | male | 176 | 85 | 73.1 | 26.9 |
| 7841 | 06-06-2023 | male | 218 | 98 | 326.8 | 146.3 |
| 7844 | 06-06-2023 | male | 181 | 90 | 400.3 | 275.0 |
| 7847 | 19-07-2023 | female | 183 | 81 | 87.5 | 70.4 |
| 7850 | 19-07-2023 | female | 152 | 73 | 52.1 | 48.3 |
| 7853 | 19-07-2023 | female | 150 | 72 | 93.4 | 31.2 |
| 7856 | 19-07-2023 | female | 160 | 80 | 60.2 | 44.0 |
| 7859 | 19-07-2023 | female | 176 | 77 | 53.6 | 67.2 |
| 7862 | 19-07-2023 | male | 140 | 70 | 42.4 | 25.0 |
| 7865 | 19-07-2023 | female | 179 | 79 | 71.2 | 35.9 |
| 7868* | 19-07-2023 | male | 128 | 66 | 1.0 | 0.0 |
| 7871 | 19-07-2023 | male | 159 | 78 | 104.6 | 89.7 |
| 7874* | 19-07-2023 | female | 199 | 86 | 25.0 | 0.0 |
| 7877 | 31-08-2023 | male | 166 | 86 | 138.6 | 141.1 |
| 7880 | 31-08-2023 | male | 157 | 80 | 110.7 | 54.4 |
| 7883 | 31-08-2023 | male | 178 | 92 | 180.0 | 150.0 |
| 7886 | 31-08-2023 | male | 175 | 86 | 14.7 | 13.3 |
| 7889 | 31-08-2023 | male | 179 | 85 | 64.1 | 145.7 |
| 7892 | 31-08-2023 | female | 157 | 74 | 41.1 | 53.6 |
| 7895 | 31-08-2023 | male | 165 | 85 | 187.5 | 13.4 |
| 7898 | 31-08-2023 | male | 161 | 83 | 83.0 | 1.0 |
| 7901 | 31-08-2023 | male | 180 | 86 | 55.5 | 48.0 |
| 7904* | 31-08-2023 | male | 175 | 85 | 0.0 | 0.0 |
| 7907 | 31-08-2023 | male | 177 | 85 | 20.1 | 17.1 |

In parallel, environmental conditions (temperature and salinity) were recorded each month by the managers of the natural reserve in the subsurface using a multiparameter probe (In-Situ AquaTROLL 500) at several stations in the lagoon (Fig. 1B).

The receiver network in the lagoon was composed of 12 receivers (models TBR 700 and TBR 800 – Thelma Biotel) (Fig 1.). The receivers were positioned on poles at specific locations to enable the detection of the movements of *C. sapidus* individuals within the lagoon, as well as potential migrations to the sea. Because *C. sapidus* individuals have the potential to access the marine environment either via the sea channel or through the Fossone channel, located near the mouth of the Golo River, three receivers were positioned along the sea channel and two along the Fossone channel. This gate□like configuration enabled detection of individuals within these channels, as well as inference of movement direction between the lagoon and the sea. Furthermore, three receivers were positioned in Forca Cove, and an additional three were situated in the central region of the lagoon, between the northern and southern basins. Ultimately, a single receiver was situated in close proximity to the Bevinco river mouth in the northwest region of the lagoon. The detection range was determined in July 2024 in similar conditions than in 2023, by means of a range test in accordance with the methodology described by Richard et al. (2020) and How & de Lestang (2012) (Supplementary Fig S2.). A transmitter was fixed to a pole in one location in the North basin of the lagoon. Receivers were placed for 7 days on poles in an aligned manner at 0 m (pole carrying the transmitter), 50 m, 100 m and 150 m. The average hourly detection rate was then computed for each distance, as well as the average detection efficiency. D50, which represents the distance at which the detection efficiency declines to 50% of the maximum detection proportion was estimated here at 100.5 meters (Supplementary Fig S2.). At sea the receiver network consisted of 58 receivers (TBR 700/800, Thelma Biotel) (Fig 1.), with an estimated detection range D50 of approximately 150 meters. Thirty-four receivers were placed between Bastia and the Biguglia lagoon (Lido de la Marana), and the remaining units were arranged into four coast□perpendicular detection barriers, each consisting of six receivers, along the eastern coast of Corsica. The receivers in the lagoon were removed on 24 October 2023, while those deployed at sea were retrieved later, on 7 December 2023.

### 2.3. Data Analysis

Detections were first sorted according to the detection frequency of the transmitter, its identifier and the date, with the objective of retaining only those detections that are linked to *C. sapidus* individuals. The raw values for depth, activity and temperature are discrete values between 0 and 255; therefore, they must be converted in order to obtain the real values. The pre-processing and data conversion were conducted using R 4.3.1 (R Core Team, 2023). The conversion was carried out by applying a coefficient, provided by the manufacturer (Thelma Biotel), in accordance with the type of variable and sensor. As the lagoon temperatures were higher than the transmitter’s detection range: [0 - 25.5 °C], these data could not be used. However, the detections linked to these values were nevertheless employed in the spatial analysis.

The use of activity data, spanning a range of 0.013588 to 0.346494 m.s□², enables the quantification of mechanical energy expenditure in small aquatic animals with a length of less than 3 meters (Martin Lopez et al., 2022). The activity calculation method employed in this study is the root mean square (RMS) method. Subsequently, this value was converted in order to ascertain the distinct activity regimes of the individuals in question, namely resting, walking and swimming (Supplementary Fig S3.). To differentiate the activity levels associated with each of the three regimes, the k-means clustering method with bootstrapping (n = 100) was used to partition the activity data into three corresponding clusters. Subsequently, a multinomial generalized additive model (GAM) was fitted using the R package ‘*mgcv’* (Wood, 2025) to investigate the effects of sex (females as reference category), diel period (day/night with day as reference category), their interaction, and the previous behavioural state on crab activity. Standardized time covariate was included as a smooth term to account for potential non-linear temporal variation, whereas crab identity was included as a random intercept effect to account for repeated observations of individuals. The previous behavioural state was included to account for temporal autocorrelation in behavioural activity. The response variable comprised three behavioural states (resting, walking, and swimming), with resting behaviour designated as the reference category. The relative frequency of each activity state was calculated as a percentage for graphical representation. For each sex, the number of detections associated with a given activity state was expressed relative to the total number of detections recorded during daytime and night-time periods. To investigate the factors influencing crab depth use, we fitted a generalized additive mixed model (GAMM) with depth as the response variable. Fixed effects included sex, diel period (Day/Night), their interaction, and crab size (width). Time was incorporated as a smooth term to account for potential non-linear temporal variation in depth use. Individual identity (crab_ID) was included as a random intercept to account for repeated observations of the same crab. Preliminary inspection of the autocorrelation function (ACF) revealed strong positive temporal autocorrelation among consecutive depth observations. Consequently, an autoregressive correlation structure of order 1 [AR(1)] was incorporated into the model to account for temporal dependence within individuals. Examination of the residual ACF after model fitting showed autocorrelation coefficients close to zero across all lags, indicating that the AR(1) structure adequately captured temporal dependence and that residuals were effectively independent. Depth data were analyzed according to the nycthemeral cycle and the sex of the individuals. Detections recorded at sea were excluded to better visualize sex-based differences in depth distribution within the lagoon.

The individual detections at each receiver are employed in the calculation of centers of activity (COAs) through the use of the ‘VTrack’ package (Campbell et al., 2012). COAs provide a two-dimensional point corresponding to the weighted mean position over a specified time interval (Kraft et al., 2023). The time step was set at 30 minutes due to the high level of observed activity exhibited by the blue crab. The cumulative distances traveled by each individual from the date of release were calculated on the basis of COAs at 60-minute intervals using the R *‘sf’* package (Pebesma, 2018). To investigate sex-specific temporal patterns in cumulative movement, cumulative distance travelled (km) was analysed using a generalized least-squares (GLS) model. Crab size (carapace width), sex, and time since release were included as explanatory variables. Because exploratory analyses indicated a non-linear relationship between cumulative distance and time, both linear and quadratic terms for time since release were incorporated into the model. To test whether temporal trajectories differed between sexes, interactions between sex and both the linear and quadratic terms of time since release were also included. The initial model included crab identity as a random intercept; however, among-individual variance was negligible and the model resulted in a singular fit. Consequently, the random effect was removed and inference was based on the GLS model. Because repeated observations collected from the same individual through time exhibited strong temporal autocorrelation, a first-order autoregressive correlation structure [AR(1)] was incorporated into the model. Residual autocorrelation was assessed using autocorrelation function (ACF) plots, and the adequacy of the AR(1) structure was verified by examining the residual temporal dependence after model fitting.

Subsequently, the COAs are employed to ascertain the extent of the home range, defined as the area traversed by an animal during its daily activities (Kraft et al., 2023). The area of occupation was determined non-parametrically using the Minimum Convex Polygon (MCP) and the Kernel Utilization Distribution (KUD) methods, with a bivariate normal fixed kernel (March et al., 2010). The MCP was employed to determine the convex hull, while the KUD was used to estimate the utilization distribution. In order to avoid overestimation of home ranges, the MCP method was employed (March et al., 2010). Minimal convex polygons were calculated for individuals detected at more than four different stations using the ‘adehabitatHR’ v.0.4.18 package (Calenge, 2020). Furthermore, KUDs are calculated for presence probabilities between 5 and 95% (in 5% steps), in order to demonstrate the heterogeneity of spatial dynamics. Subsequently, the 50% and 95% KUDs are isolated, as they correspond to the core area of the home range and the global home range, respectively. The smoothing parameter employed is the reference parameter, designated as “href.” This method was applied to individuals with a minimum of 30 detections during the study period. Subsequently, the KUDs were processed in QGIS 3.34, due to the lagoon’s distinctive morphology. Accordingly, the KUDs were trimmed to align with the topography of the site, thus preventing an overestimation of the areas in question. This was achieved by incorporating data from adjacent lands or, in select instances, from marine sources, where no detections were documented. In the case of KUD layers that overlap with the San Damiano peninsula, the portion of the KUD corresponding to the percentage of the layer in question was removed from Forca Cove. Furthermore, the ad hoc method was employed to prevent the overestimation of KUDs for individuals detected at sea. In order to isolate the polygons at sea and add them to the KUDs in the lagoon, different smoothing parameters were applied. These manipulations are conducted with the objective of obtaining realistic estimates of the areas of occupation of *C. sapidus* in the lagoon and at sea. Subsequently, the polygons were processed and analyzed using R 4.3.1 (R Core Team, 2023). Differences in KUD 50% and KUD 95% between male and female blue crabs were assessed using non-parametric Wilcoxon rank-sum tests.

The aforementioned measurements were employed in two principal analytical procedures. Firstly, a kriging interpolation was performed in order to generate spatial representations of salinity, which were then linked to the monthly detection proportions of each sex. Secondly, the occurrence frequencies across the study period were assessed based on salinity and temperature ranges. A Quotient Index (QI) was calculated by dividing the frequency of occurrences in each range by the proportion of stations in those salinity ranges, multiplied by 100. Furthermore, detections occurring at sea were analyzed in conjunction with average monthly salinity and temperature data, sourced from the Copernicus Marine Environment Monitoring Service.

## 3. Results

### 3.1. Temporal survey

Of the 31 tagged crabs (11 females and 20 males), one transmitter (ID 7904) was never detected. All other transmitters were detected at least once during the study (Table 1, Figure 2.). A total of 62, 580 detections were recorded, with 11, 360 for females and 51 220 for males. The mean number of detections per individual was 2 086, with a range from one detection for transmitter 7868 to 17 159 detections for transmitter 7844. The mean detection period per individual was 35 days, with a range from a few hours (ID 7835) to 129 days, over four months (ID 7844). The cessation of detection could be attributed to one of several factors, including movement outside the receivers’ network, moulting, mortality, transmitter loss, or the animal’s capture. Five individuals were excluded from the analyses: two due to either a lack of detections or single detections, and three that only recorded activity on the day of release or showed consistently low activity at the same depth as indicative of mortality or transmitter detachment.

**Figure 2.**
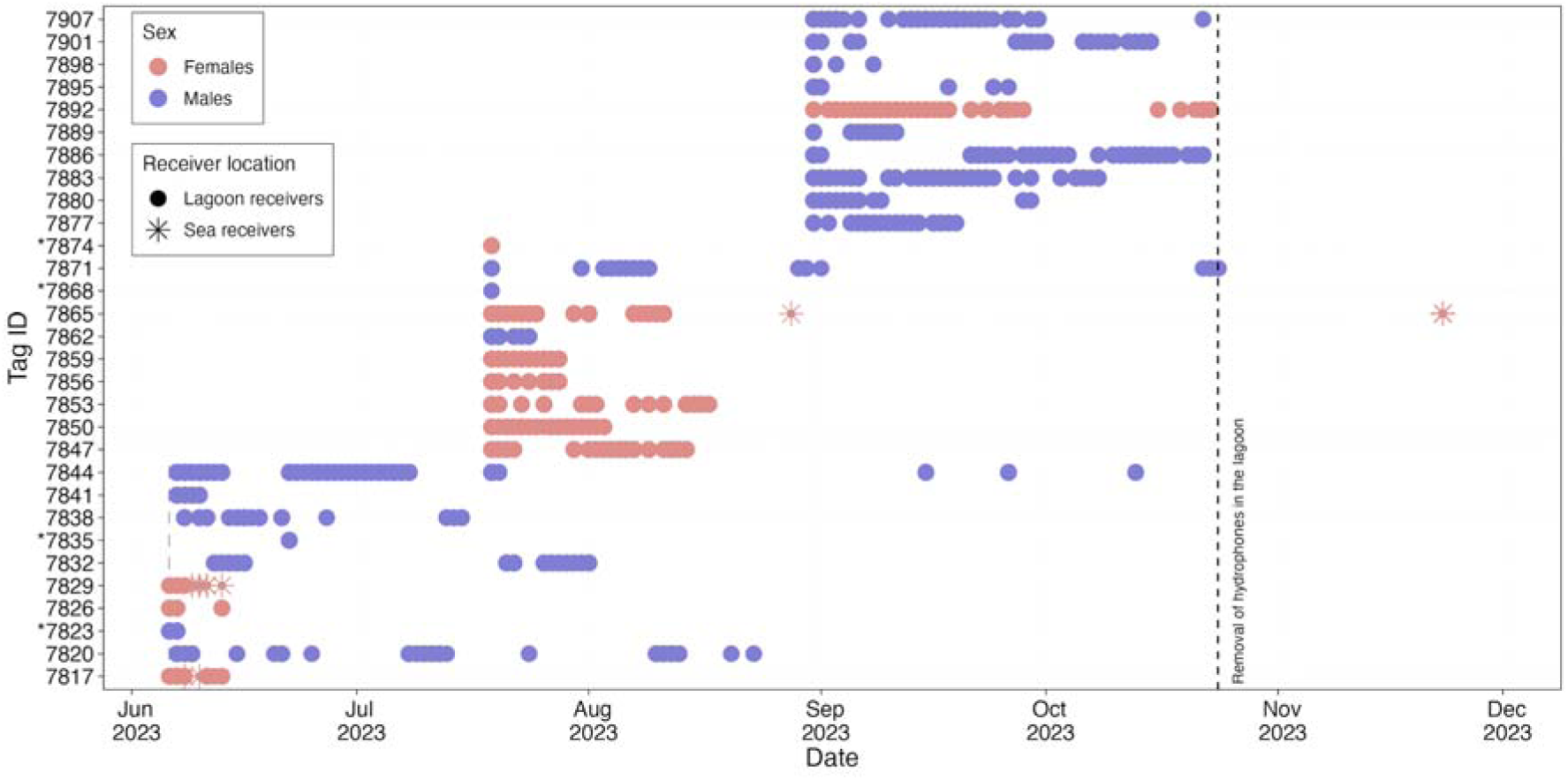
Abacus plot of individual detections of the blue crab (*Callinectes sapidus*) recorded during acoustic telemetry survey in the Biguglia lagoon in 2023. The dotted line corresponds to the removal of the hydrophones in the lagoon on 24th of October 2023. Note: Individuals marked with an asterisk (*) were excluded from the analysis due to one or more of the following reasons: absence of detections, single detection events, activity recorded only on the day of release, or consistently low activity at a fixed depth.

During the study period, three female individuals were identified by receivers at the sea (Figure 2). Transmitters 7817 and 7829 were detected between the 8th and 10th of June near the sea channel, on the sea receiver located north of the sea channel and closest to the shoreline. Transmitter 7817 was then detected in the lagoon between the 12th and 14th of June. Transmitter 7865 was first detected at sea on 28 August by the receiver located near the mouth of the Golo River, and again on 23 November. This individual was subsequently recovered by a fisher in December with its transmitter near Moriani Beach, approximately 15 km south of the last detection point.

### 3.2. Activity and depth distribution

Overall, resting was the predominant behavioural state, accounting for a mean (± SD) of 64.9 ± 16.2% of observations during the day and 62.3 ± 24.0% at night. Walking was the second most common activity (28.8 ± 10.6% during the day and 31.6 ± 16.6% at night), whereas swimming was the least frequent behavioural state, representing 11.9 ± 19.7% and 12.7 ± 19.9% of observations during the day and night, respectively (Figure 3; Supplementary Fig. S3). Marked differences were observed between sexes. Females were generally more active than males, exhibiting lower proportions of resting behaviour and higher proportions of locomotor activities. During the day, females spent 54.9 ± 15.8% of observations resting, 33.0 ± 10.5% walking and 20.9 ± 28.4% swimming, whereas males spent 70.6 ± 14.0% of observations resting, 26.2 ± 10.1% walking and only 5.5 ± 4.4% swimming. At night, female resting behaviour decreased further to 48.3 ± 23.0%, accompanied by an increase in walking activity to 39.9 ± 17.0%, while males remained predominantly inactive (71.6 ± 20.4% resting). These patterns indicate greater nocturnal activity in females and a predominance of resting behaviour in males.

**Figure 3.**
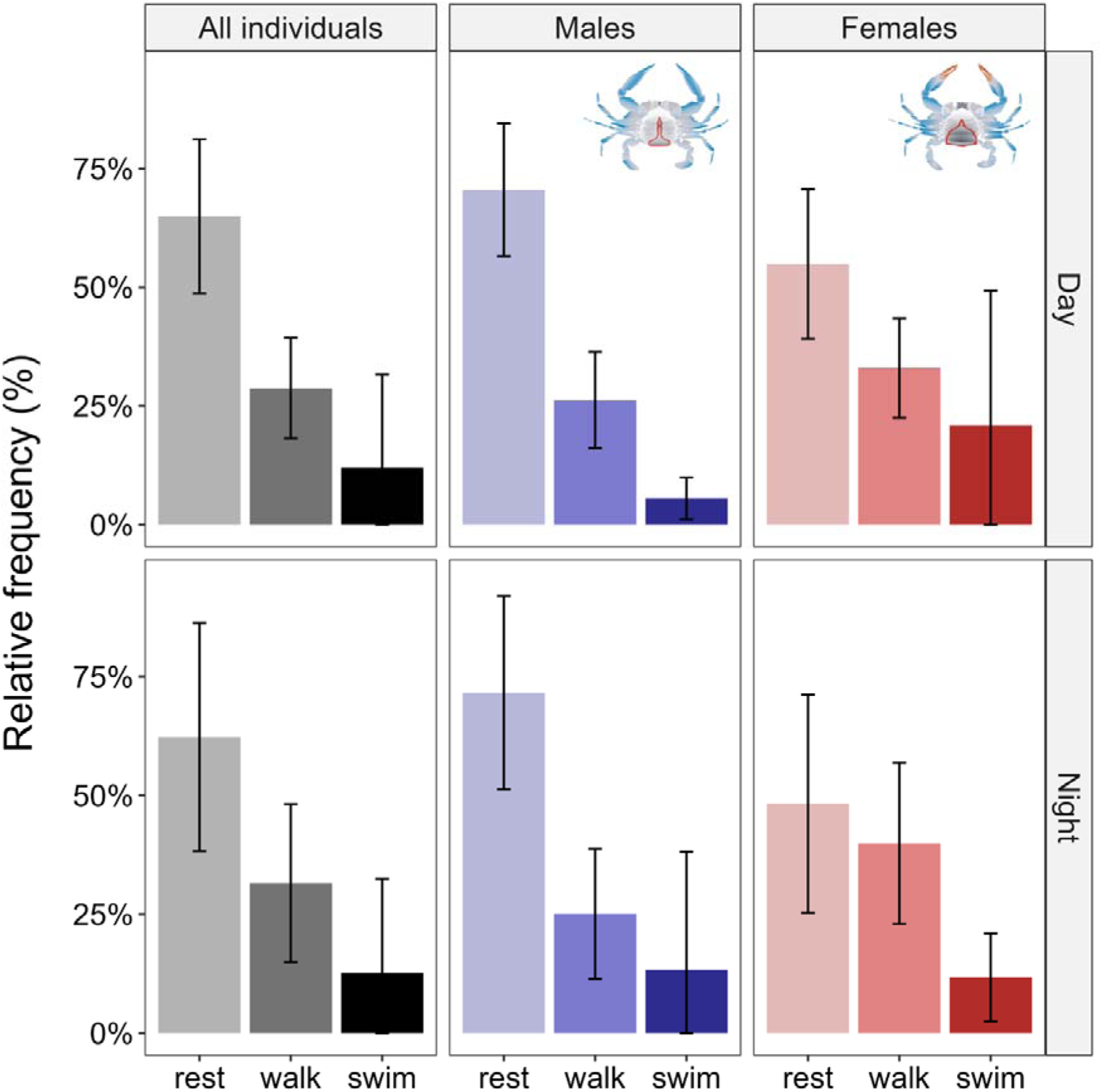
Activity patterns of the blue crab (*Callinectes sapidus*) recorded during acoustic telemetry survey in the Biguglia lagoon in 2023. Bars represent the mean (± SD) relative frequencies of resting, walking, and swimming behaviours during daytime and night-time periods.

The multinomial GAM assessing the effects of previous behavioural state, sex, diel period, their interaction, temporal variation, and individual variability on crab activity explained 24.4% of the deviance and was based on 20,831 observations from 26 individuals (Table 2). Previous behavioural state was the strongest predictor of current activity. Crabs that were swimming at the previous observation were substantially more likely to be swimming than resting (β = 3.58, SE = 0.10, p < 0.001), while crabs that were previously walking were more likely to be walking than resting (β = 2.61, SE = 0.04, p < 0.001). Similarly, crabs that were previously walking were more likely to swim than rest (β = 2.49, SE = 0.08, p < 0.001), and crabs that were previously swimming were more likely to walk than rest (β = 2.59, SE = 0.08, p < 0.001), indicating strong behavioural persistence through time.

**Table 2.** Results of the multinomial generalized additive model (GAM) assessing the effects of previous activity state, sex, diel period (day/night), their interaction, standardized time, and individual variability on behavioural activity (‘swim’ and ‘walk’, ‘rest’ as reference category) of the blue crabs (*Callinectes sapidus*) recorded during acoustic telemetry survey in the Biguglia lagoon in 2023. Parametric coefficients, smooth-term statistics, and overall model fit metrics are presented. Abbreviations: SE, Standard Error; edf, Effective Degrees of Freedom; χ², Approximate Chi-square Statistic.

| Contrast | Variable | Estimate | SE | z | p-value |
| --- | --- | --- | --- | --- | --- |
| 'swim' vs 'rest' | (Intercept) | -3.439 | 0.238 | -14.45 | <0.001*** |
|  | activity_lag_swim | 3.579 | 0.105 | 34.18 | <0.001*** |
|  | activity_lag_walk | 2.493 | 0.081 | 30.88 | <0.001*** |
|  | Sex (ref. = female) | -0.662 | 0.301 | -2.20 | 0.028* |
|  | Day_Night (ref. = Day) | 0.002 | 0.133 | 0.02 | 0.987 |
|  | Sex × Day_Night | -0.082 | 0.158 | -0.52 | 0.605 |
| 'walk vs 'rest' | (Intercept) | -1.904 | 0.155 | -12.32 | <0.001*** |
|  | activity_lags_wim | 2.588 | 0.081 | 31.92 | <0.001*** |
|  | activity_lag_walk | 2.606 | 0.042 | 62.75 | <0.001*** |
|  | Sex (ref. = female) | -0.215 | 0.197 | -1.09 | 0.275 |
|  | Day_Night (ref. = Day) | 0.255 | 0.092 | 2.79 | 0.005** |
|  | Sex × Day_Night | -0.486 | 0.102 | -4.75 | <0.001*** |
|  | **Smooth term** | **edf** | **χ<sup>2</sup>** | **p-value** |  |
| 'swim' vs 'rest' | s(time_sc) | 7.53 | 78.25 | <0.001*** |  |
|  | s(ID_crab) | 18.47 | 254.31 | <0.001*** |  |
| 'walk vs 'rest' | s(time_sc) | 7.44 | 50.66 | <0.001*** |  |
|  | s(ID_crab) | 19.07 | 233.21 | <0.001*** |  |
|  | **Indicator** | **Value** |  |  |  |
|  | N individuals | 26 |  |  |  |
|  | n observations | 20 831 |  |  |  |
|  | Deviance explained | 24.4 % |  |  |  |

Consistent with the activity budget analysis (Figure 3), males were significantly less likely to swim than females (β = −0.662, SE = 0.301, p = 0.028). No significant effect of diel period was detected on swimming activity (β = 0.002, SE = 0.133, p = 0.987), and the interaction between sex and diel period was not significant (β = −0.082, SE = 0.158, p = 0.605). For the walk-versus-rest contrast, males did not differ significantly from females (β = −0.215, SE = 0.197, p = 0.275). However, crabs were more likely to walk at night (β = 0.255, SE = 0.092, p = 0.005), and a significant Sex × Day_Night interaction (β = −0.486, SE = 0.102, p < 0.001) confirmed that nocturnal changes in activity differed between sexes, with females exhibiting a greater increase in walking activity at night.

Temporal effects were highly significant for both multinomial logits (swim vs. rest: edf = 7.53, χ² = 78.25, p < 0.001; walk vs. rest: edf = 7.44, χ² = 50.66, p < 0.001), indicating non-linear variation in behavioural activity through time. Significant individual effects were also detected for both contrasts (swim vs. rest: edf = 18.47, χ² = 254.31, p < 0.001; walk vs. rest: edf = 19.07, χ² = 233.21, p < 0.001), revealing substantial among-individual heterogeneity in behavioural patterns.

Focusing solely on lagoon detections, depth use was significantly affected by diel period, with crabs occupying slightly shallower depths at night than during the day (β = −0.024 ± 0.010 SE, p = 0.021; Supplementary Fig S4., Table S1.). A significant non-linear temporal effect was also detected (edf = 8.69, F = 24.67, p < 0.001). Individual variation in depth use was substantial (random intercept SD = 0.347 m), and an AR(1) structure adequately accounted for the strong temporal autocorrelation observed in the data (φ = 0.913). No significant effects of sex or crab size on depth use were detected (Supplementary Table S1.). Males occupied a mean depth of 1.56 ± 0.36 m during the day and 1.50 ± 0.41 m at night. Within the lagoon, the maximum depth recorded for females was 2.6 m, while males reached up to 3.1 m. Females that migrated to the sea were detected at depths of up to 6.8 m near the sea channel and as deep as 9.3 m at the mouth of the Golo River.

### 3.3. Area utilization analysis

#### 3.3.1. Distances

Individual cumulative traveled distance ranged from 0.0 to 6.8 km by day 10, 1.0 to 18.9 km by day 25, and 2.2 to 24.2 km by day 50. Correspondingly, average individual hourly speeds varied from 0.0 to 28.3 m.h^-1^ on day 10, between 1.7 to 31.5 m.h^-1^ on day 25, and 3.6 to 40.3 m.h^-1^ at day 50.

Cumulative distance travelled increased significantly with time (β = 0.338 ± 0.018 SE, p < 0.001), although the significant negative quadratic effect of time (β = −0.0020 ± 0.0001 SE, p < 0.001) indicated a progressive reduction in the rate of distance accumulation through time (Table 3; Fig. 4). Neither sex nor crab size significantly affected cumulative distance travelled. However, significant interactions between sex and both the linear (β = −0.228 ± 0.020 SE, p < 0.001) and quadratic (β = 0.0013 ± 0.0001 SE, p < 0.001) effects of time revealed distinct temporal trajectories between sexes. Females accumulated distance more rapidly and reached greater cumulative distances over the monitoring period, whereas the rate of distance accumulation in males declined more rapidly, resulting in an earlier plateau (Fig. 4). Temporal autocorrelation between successive observations was high and was accounted for by the AR(1) correlation structure (φ = 0.997). A high degree of individual variability was observed, with some individuals traveling very long distances while others remain relatively sedentary. Additionally, several trajectories show a plateau in movement, reflecting the intermittent detection by a single receiver.

**Figure 4.**
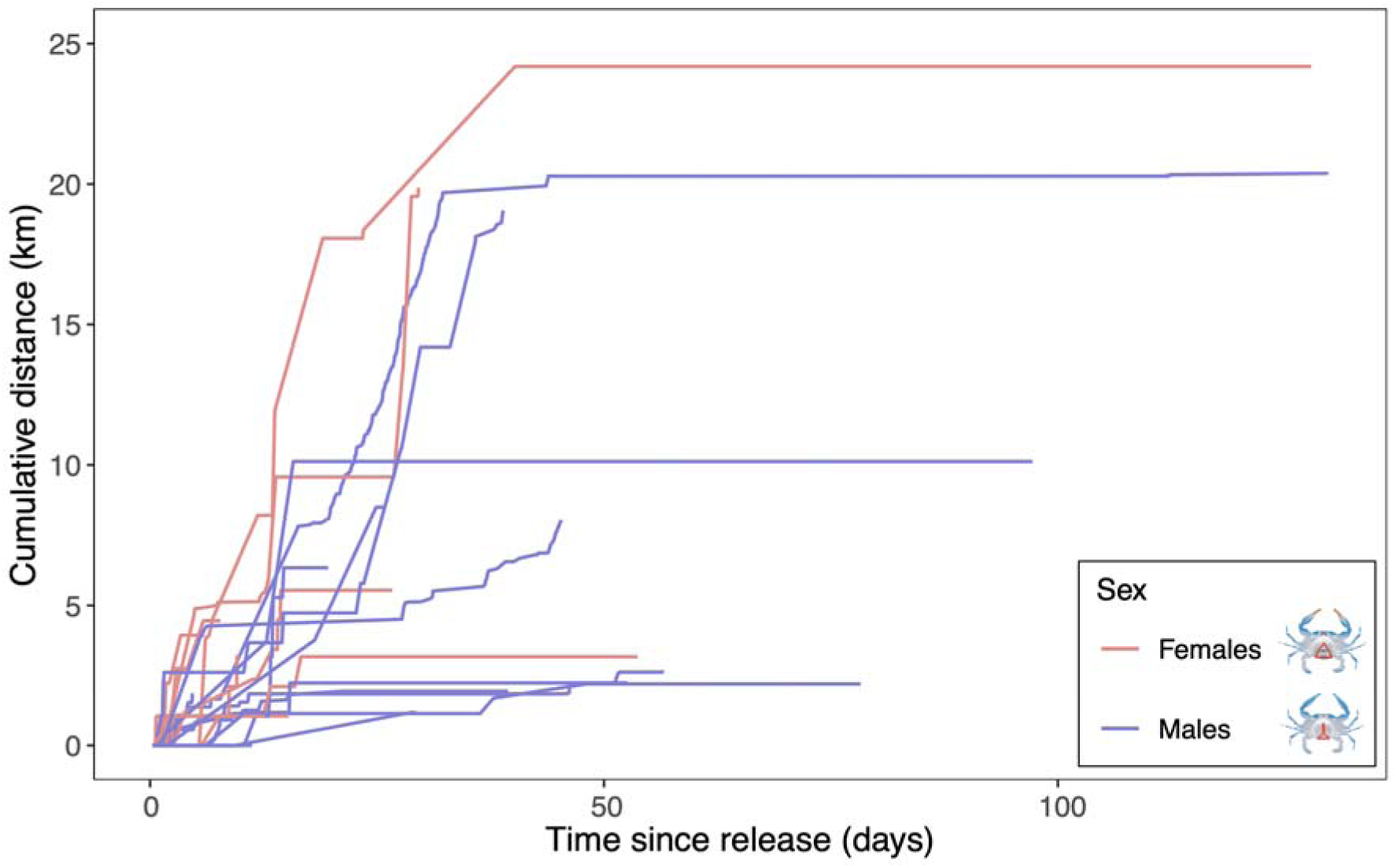
Cumulative distance travelled by each individual of the blue crab (*Callinectes sapidus*) recorded during acoustic telemetry survey in the Biguglia lagoon in 2023. A distinction is made between females (red lines) and males (blue lines).

**Table 3.** Results of the generalized least-squares (GLS) model examining the effects of sex, crab size and temporal changes in cumulative distance travelled by the blue crabs (*Callinectes sapidus*) recorded during acoustic telemetry survey in the Biguglia lagoon in 2023. The model included linear and quadratic effects of time since release and their interactions with sex. Parametric coefficients, autocorrelation structure, and overall model fit metrics are presented. Abbreviations: SE, Standard Error; AR(1), first-order autoregressive temporal correlation structure; φ, autoregressive parameter. NB1: The individual random effect (ID_crab) was removed from the final model because it explained negligible variance and resulted in a singular fit.

| **Variable** | **Estimate** | **SE** | **t-value** | **p-value** |
| --- | --- | --- | --- | --- |
| Intercept | -3.559 | 9.791 | -0.36 | 0.716 |
| Sex (ref. = female) | 1.077 | 1.998 | 0.54 | 0.590 |
| Crab size (mm) | 0.025 | 0.059 | 0.42 | 0.676 |
| Time | 0.338 | 0.018 | 18.60 | <0.001*** |
| Time<sup>2</sup> | -0.0020 | 0.0001 | -17.70 | <0.001*** |
| Sex × Time | -0.228 | 0.020 | -11.42 | <0.001*** |
| Sex × Time<sup>2</sup> | 0.0013 | 0.0001 | 9.44 | <0.001*** |
| **Autocorrelation structure** |  |  |  |  |
| AR(1) parameter ($\phi$) | 0.997 | | | |
| **Indicator** | **Value** |  |  |  |
| N individuals | 25 |  |  |  |
| n observations | 2 321 |  |  |  |
| Adjusted R<sup>2</sup> | 21.0% |  |  |  |

#### 3.3.2. Kernel Utilization Distribution

The home-range sizes were calculated using KUDs for a total of 17 individuals, 8 females and 9 males. In general, females tended to have larger KUD than males during the study period. Although females exhibited a larger mean KUD95 and KUD 50 areas (5.24 ± 18.40 km² and 1.65 ± 4.34 km2) than males (1.65 ± 1.70 km² and 0.42 ± 0.66 km²), the large inter-individual variability resulted in substantial overlap between sexes and no significant difference was detected (Wilcoxon rank-sum tests, p > 0.05). Two females in particular stand out by using a higher area than others, females 7853 (KUD 95 = 13.85 km²) and 7865 (KUD 95 = 23.75 km²) (Figure 5B, Figure 6). Overall KUD 95 ranged from 0.5 to 23.75 km² for females and from 0.25 to 3.35 km² for males; KUD 50 ranged from 0.20 to 5.95 km² for females and 0.10 to 1.10 km² for males.

**Figure 5.**
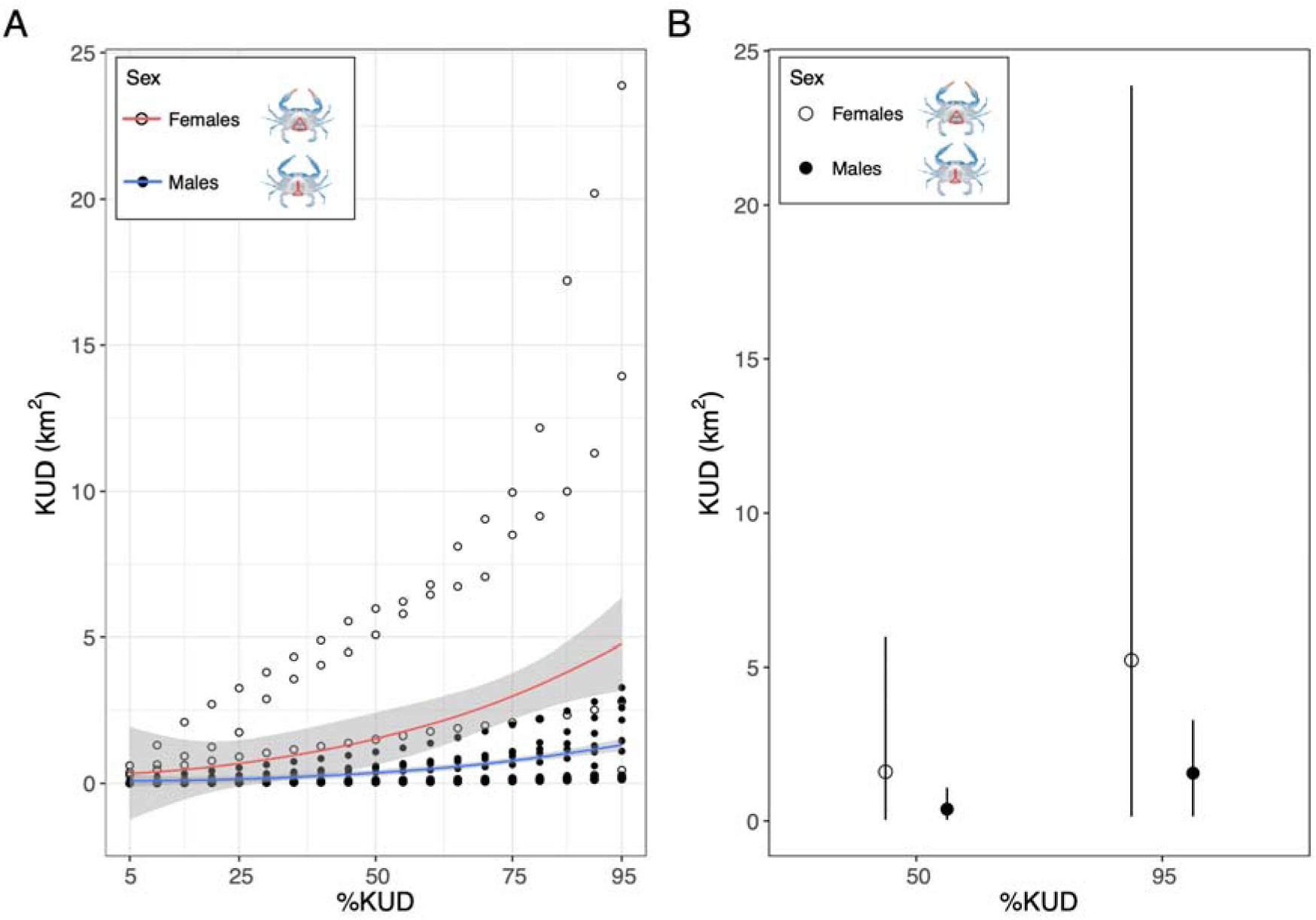
Differences in home-range size estimated using Kernel Utilization Distributions (KUDs; km²) between male and female blue crabs (*Callinectes sapidus*) monitored by acoustic telemetry in the Biguglia Lagoon during 2023. (A) KUD area (km²) across KUD levels (5–95%). Solid lines represent fitted trends for females (open circles, red line) and males (filled circles, blue line), with shaded areas indicating 95% confidence intervals. (B) Mean KUD50 and KUD95 values (± SD) for females and males.

**Figure 6.**
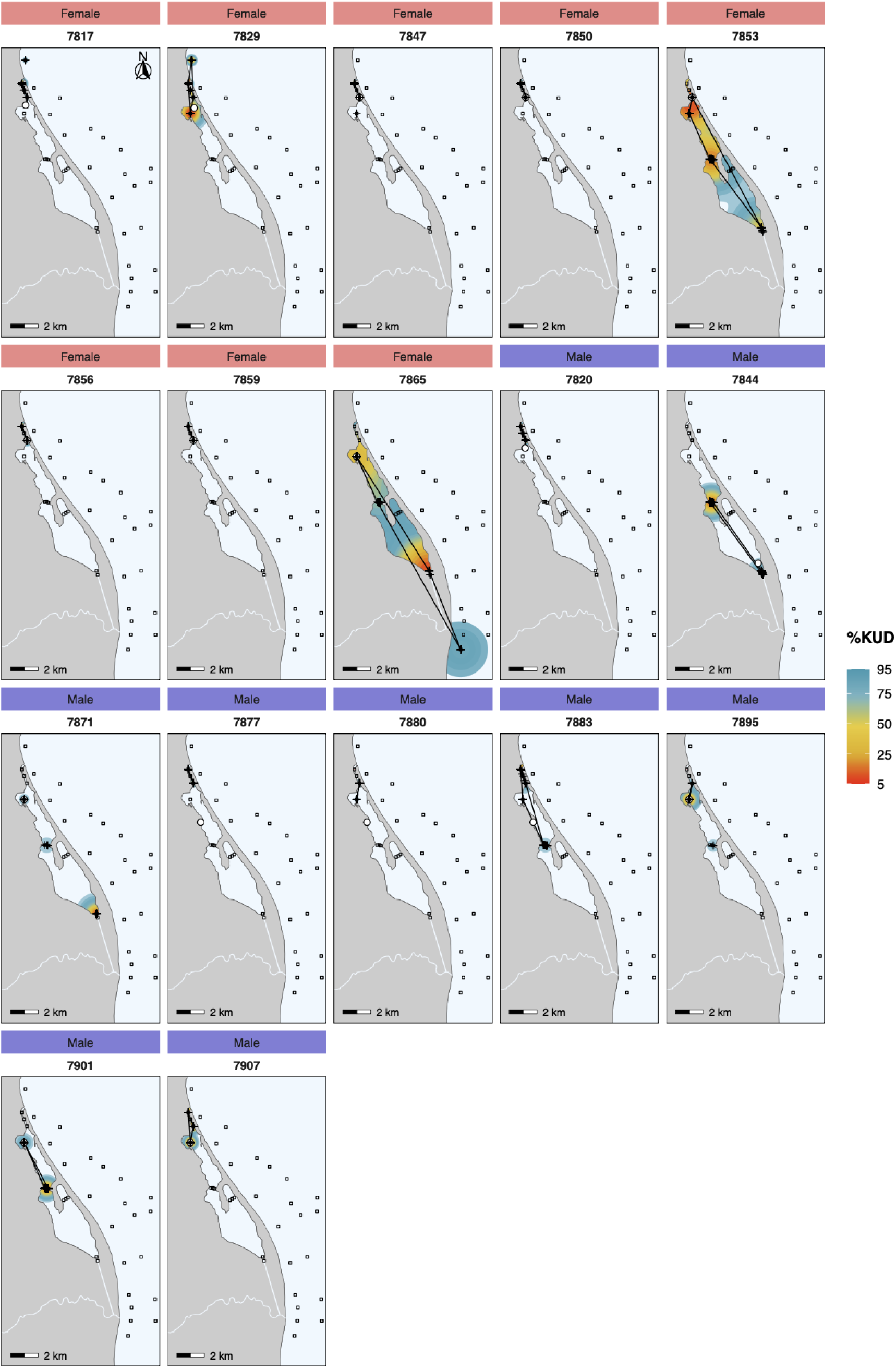
Individual home ranges of the blue crab (*Callinectes sapidus*) recorded during acoustic telemetry survey in the Biguglia lagoon in 2023. Home ranges (KUD) from 5% (red) to 95% (lightest blue), in 5% steps and minimum convex polygons (95% MCP) represented by polygons with black outlines, for each individual that could be calculated. The individual’s release site is indicated by a white dot. The stations of the two hydrophone networks are indicated by black squares, and the COAs by crosses.

Individual KUDs showed some spatial variability in habitat use between individuals (Figure 6). Although most individuals showed a localized home range, some individuals stand out with a wider home range coverage. Females 7853 and 7865 have a global home range (95% KUD) covering almost the entire lagoon and even beyond for female 7865, having part of the home range at sea to the south of the lagoon at the level of the Fossone channel. Males tend to occupy more of the southern part of the northern basin, while females tend to occupy the sea channel mainly.

### 3.4. Distribution of the blue crabs versus salinity

The proportion of detections at receivers in the lagoon and at sea showed a certain temporal pattern according to sex (Figure 7). Due to the removal of the marine hydrophones on 24th of October, only detections at sea could be considered in November. The females tended to occupy mainly the north part of sea channel in June, with salinities of around 15 PSU. In August, some individuals moved toward the southern part of the lagoon, corresponding to higher salinity levels ranging from 25 to 30 PSU. From September to October only a single female (ID□7892) was detected exclusively in the sea channel where salinity approached 30□PSU (Figures□2 and□7). In the case of males, in June and July they were present mainly in the northern and southern parts of the southern basin, with salinities close to 15 PSU. During August and September, male moved towards the north, with a salinity level of around 20 PSU, to be detected almost exclusively in the northern basin from September onwards with salinities between 20 and 30.

**Figure 7.**
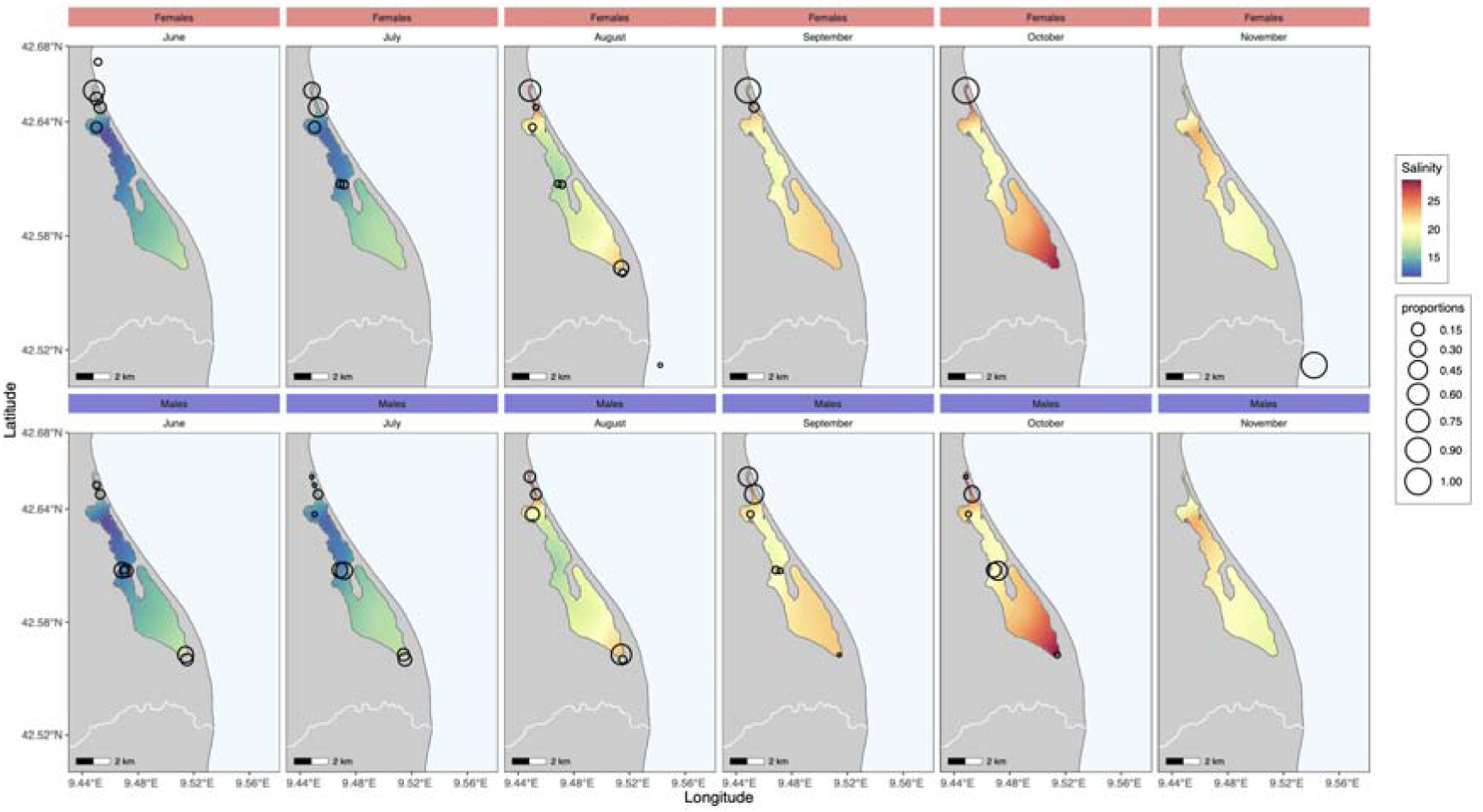
Proportion of monthly individual detections of the blue crab (*Callinectes sapidus*) at receivers for each sex associated with variation in salinity gradients recorded during acoustic telemetry survey in the Biguglia lagoon in 2023. NB: receivers in the lagoon were removed on the 24th of October.

Analysis of frequency of occurrence as a function of salinity and Quotient Index (QI) showed a pattern of habitat use across the salinity gradient (Figure 8). Females showed a bimodal preference, with QI greater than 1 at low salinity ranges (14 to 16 PSU) and higher salinity ranges (24 to 26 PSU), indicating a preference for these environments. Males, on the other hand, showed a QI>1 only for the 24-26 PSU range. Notably, a substantial proportion of males were detected at very low salinities (10 to 12 PSU), where females were rarely detected. Conversely, some females were detected at very high salinities (30 to 40 PSU), unlike males, which were not detected at such high salinities.

**Figure 8.**
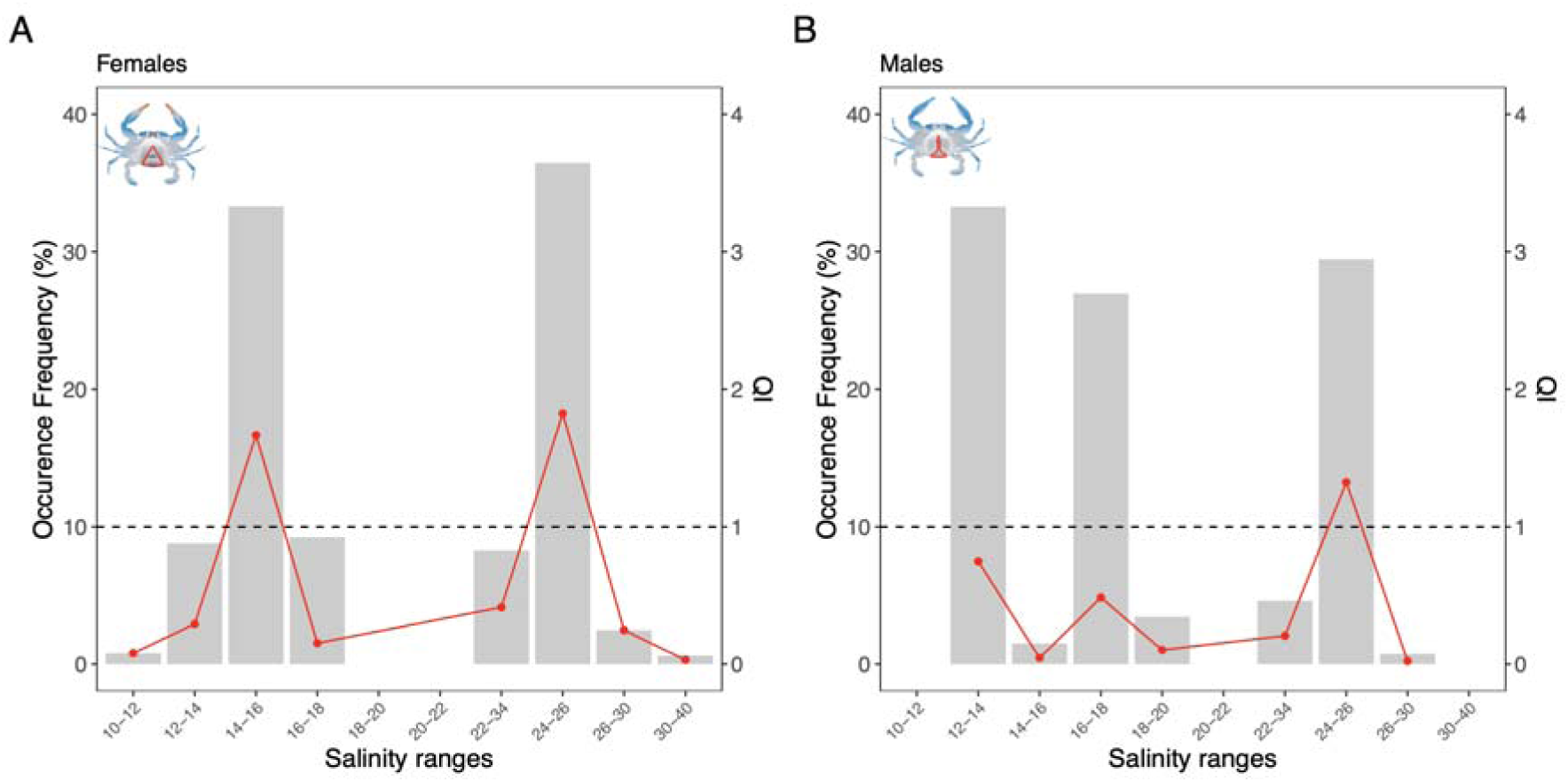
Frequency of occurrence and Quotient Index (QI>1 represents a preference) of the blue crab (*Callinectes sapidus*) by salinity range for females (A) and males (B) during the whole study period of the acoustic telemetry survey in the Biguglia lagoon in 2023.

## 4. Discussion

### 4.1. Behavioural adaptation to a novel environment

Over the past decades, and especially in the fast few years, the Atlantic blue crab (*Callinectes sapidus*) has undergone a rapid and widespread expansion throughout the Mediterranean Sea, establishing itself as a highly invasive species with significant ecological and economic consequences (Mancinelli et al., 2021; Clavero et al., 2022; Marchessaux et al., 2024b). In its native western Atlantic range, the blue crab inhabits dynamic coastal environments like estuaries and bays, characterized by strong salinity gradients and significant tidal fluctuations (Hines, 2007; Epifanio, 2019). It can also tolerate saline embayments (Ramach et al., 2009) to hypersaline conditions, such as those found in lagoons in Texas and Louisiana (Guerin & Stickle 1992), although these habitats are less studied. In contrast, its invasive range in the Mediterranean features minimal tidal variation (Ibañez et al., 2000; Sannino et al., 2015) and generally salinity gradients of lower spatial extent, especially in coastal lagoons. Given that blue crab behaviour is highly influenced by environmental conditions, it is essential to understand how the species adapts to the specific characteristics of Mediterranean lagoon systems. This study presents, to our knowledge, the first reported behavioural analysis of *C. sapidus* within its invasive range in a typical Mediterranean lagoon, Biguglia Lagoon in Corsica, using acoustic telemetry. While our findings confirm several behavioural traits previously observed in the species’ native range, they also reveal distinct behavioural adaptations specific to the Mediterranean context. Despite some characteristic high interindividual variability, we identified consistent sex-based behavioural patterns, offering valuable insights for the development of targeted management strategies.

### 4.2. Methodological Considerations

Despite the advantages of acoustic telemetry, limitations such as restricted receiver coverage and environmental interference (e.g., turbidity, acoustic noise) may have influenced detection rates (Hussey et al., 2015; Crossin et al., 2017; Ducos et al., 2022). The interpretation of the data obtained with acoustic telemetry is complex due to the influence of multiple factors, ranging from environmental conditions to technical and operational constraints that can introduce variability and biases (Huveneers et al., 2016). Nevertheless, the probability of detection relative to distance aligns with expectations for this type of environment (Richard et al., 2020). A sex bias in tagging—favoring males due to capture rates—resulted in a higher detection frequency for males. Nonetheless, the average detection period of 35 days is notable for this species, especially considering molting cycles in decapods (Guerra-Castro et al. 2011; Florko et al., 2021). Regarding the use of acceleration as a proxy of activity, there is a broad consensus about its utility in both the aquatic and terrestrial environments (Wilson et al., 2020). Although built-in accelerometers in acoustic transmitters have certain limitations for capturing fine-scale activity patterns, they are generally well suited for assessing overall activity levels (Pereñiguez et al., 2022). Finally, although spawning begins during the summer months in Mediterranean lagoons, no berried blue crab females were observed at the time of capture and tagging between early June and late August. Only one female was caught in August, and it was not carrying eggs. It is worth noting that in 2023, the year of the study, the spawning period may have preferentially occurred from late August into September.

### 4.3. Diel activity patterns and distance travelled

In our study, blue crabs exhibited activity during both daytime and night-time hours, regardless of sex, consistent with previous observations of the species in its native range (Clark et al., 1999; Bell et al., 2003; Hench et al., 2004; Darnell et al., 2012). Regarding walking activity, males were slightly more active during the day, whereas females were significantly more active at night. We did not detect different swimming activity between day and night, as it has been previously described particularly for females (Hench et al 2004, Darnell et al 2012). However, Carr et al (2004) showed that specifically during spawning migration ovigerous female crabs activity percentage was higher at night than during the day. More complexly in their macrotidal native range, female blue crabs exhibit circatidal rhythms, characterized by vertical swimming movements closely synchronized with ebb tides—a behaviour known as selective tidal stream transport—which facilitates their spawning migrations (e.g., Carr et al., 2004; Darnell et al., 2012). Movements are also highly episodic, with bursts of locomotion interspersed with rest described “swimming bouts” (Forward et al., 2005; Carr et al., 2004). Blue crabs travelled hourly mean distances ranging from 1.7 to 31.5 m.h^-1^ over a 25-day period, underscoring the high degree of interindividual variability in their movement and dispersion behaviour. This is consistent with native range studies reporting considerable variations in mean hourly speeds ranging of 0.7 to 93.2 m.h^-1^ (e.g., Hines et al., 1995; Clark et al., 1999; Bell et al., 2003). This variability is supposed to be especially high in days prior to moult (Wolcott and Hines 1990). Brief episodes of rapid, directionally oriented movements exceeding 50 m·h□¹ were also observed, consistent with patterns previously documented in the native range of the species (Wolcott & Hines, 1990; Hines et al., 1995). Overall females demonstrated higher activity walking and swimming levels and travelled greater distances, likely reflecting reproductive needs and exploratory behaviour. These behavioural differences may reflect physiological or energetic needs, with females being more active due to their reproductive role and the need to locate optimal spawning sites at sea (Carr et al 2004, Hench et al 2004, Darnell et al 2012). During spawning migrations, Darnell et al. (2012) showed that mean daily swimming activity increases from approximately 2% of the day seven days before larval release to 25% on the day of release. Although the recent bathymetry of the Biguglia lagoon is not well documented (Dufresnes et al. 2017), females were generally detected closer to the surface than males. This suggests that males may spend more time on or buried in the substrate, while females remain more active in the water column especially at night. These findings are consistent with higher activity levels and greater dispersal observed in females, likely due to swimming over longer distances. Our results finally demonstrate that blue crabs especially females are capable of sustained swimming activity and long-distance travel even in the absence of a strong tidal regime, highlighting their remarkable behavioural plasticity in Mediterranean coastal lagoon environments.

### 4.4. Spatial Distribution and Habitat Use

Spatial analysis also revealed sex-based differences in habitat use. Males occupied a broader range of lagoon zones, including lower-salinity areas, while females favoured higher-salinity regions and exhibited larger home ranges. KUD analysis revealed substantial interindividual variability in spatial use patterns, consistent with previous findings, along with marked differences between males and females. This behaviour suggests a broader and more dynamic habitat use by females likely linked to reproductive strategies. In their native range, blue crab mating typically occurs in upper estuaries after the female’s terminal moult (Ramach et al., 2009). Males display some specific stationary paddling courtship behaviour to attract females (Wood & Derby 1995, Kamio et al 2008). An initial sex-based difference in habitat use may reflect female avoidance behaviour in areas of high male density, a pattern consistent with the agonistic interactions and resulting sex segregation commonly observed in blue crabs outside the mating period (Hines, 2007). The sex ratio in the Biguglia lagoon is skewed toward males (Marchessaux et al., 2024b) potentially prompting females to adopt such behaviour. Post-mating, females forage to restore energy and allow ovarian maturation (Aguilar et al., 2005). Females then migrate seaward to spawn, releasing larvae in marine environments (Hines et al., 2003; Darnell et al., 2009). Embryonic development lasts about 2–3 weeks before hatching (Epifanio 2019). Females produce multiple clutches, often returning to estuarine areas between releases (Hench et al., 2004). Over time, spawning females accumulate in high-salinity waters, enhancing larval dispersal success (Forward et al., 2005; Rittschof et al. 2010). In our study, three females exhibited extensive spatial use, with home ranges encompassing the entire lagoon and extending into adjacent marine areas. These individuals were detected at sea: two migrated through the channel in June—one of which returned to the lagoon shortly after—and one exited via the Fossone channel in the south in August and remaining at sea. Notably, all three migrations occurred within days of tagging, and none of the females were ovigerous at the time. Only the female that exited via the Golo estuary more than a month after release could have had sufficient time to produce and release larvae at sea. We also observed that the sea channel of the Biguglia Lagoon experienced intermittent closure events due to sand accumulation, lasting from several days to over a week during the study period (unpublished data). This temporary open–closed configuration is characteristic of Mediterranean lagoonal and estuarine systems (Erostate et al., 2022). Similarly, the Golo Estuary undergoes regular closure events, particularly during the summer months, primarily due to reduced river discharge associated with dam operations for hydroelectric power generation. These closure events may significantly limit connectivity between the lagoon and the sea, potentially restricting the movement of female blue crabs and leading to increased female densities in lagoon areas near the openings. Despite these potential barriers, blue crabs have been observed traversing emerged sandy areas in other Mediterranean locations (Marchessaux, pers. comm.). These observations indicate that both non-ovigerous and potentially ovigerous females are capable of undertaking early seaward migrations, for exploratory purposes or to complete reproductive maturation in marine environments.

As a successful euryhaline species, the blue crab is capable of thriving in both brackish and hypersaline environments within its native range. This remarkable adaptability is supported by localized physiological mechanisms (Guerin & Stickle, 1992). In this study, male blue crabs exhibited a clear preference for lower- to intermediate-salinity environments, gradually shifting toward these specific areas of the lagoon over time. While females were also present in intermediate-salinity zones, they progressively moved toward higher-salinity areas beginning in August, the period of the main spawning migration for the studied year. When available, blue crabs tend to prefer these low to intermediate salinity levels, which are typically found in estuarine environments (Hines, 2007). This preference for intermediate salinities helps them maintain optimal internal conditions and reduces the stress on their osmoregulatory mechanisms (Tagatz, 1971). Lower salinity water is also associated to higher food production and availability which might provide a nutritional advantage (Peterson, 2003). Salinity also influences aerobic metabolism and growth (Guerin & Stickle 1992, Marchessaux et al 2024a). The results of this Mediterranean study align with existing literature, confirming salinity as a critical environmental factor affecting growth, survival, behaviour, reproduction, and distribution (Aguilar et al., 2005; Ramach et al., 2009; Marchessaux et al. 2024a,b).

### 4.5. Implications for management

This study has directly informed the development of a Territorial Control Plan for the Atlantic blue crab (*Callinectes sapidus*) in Corsica. By the end of 2023, drawing upon a comprehensive body of scientific research and field data (Marchessaux et al. 2024 a,b), a dedicated management strategy was drafted and subsequently approved unanimously by the Corsican Assembly in December 2024. This marks the first such initiative in France targeting a marine invasive species. The Control Plan serves as a regional adaptation of the national strategy, adapted to the specific Corsica’s unique ecological and socio-economic context. Its overarching objective is to provide a structured framework based on the five pillars of the national strategy, aimed at strengthening coordination, organizing collective efforts, and implementing targeted actions grounded in robust scientific evidence. It has become clear that eradication — the complete removal of all individuals from an invaded area — is not a feasible option for the blue crab, whose populations are now viable, reproducing, and integrated into local ecosystems (UNEP/MAP-SPA/RAC 2025). The challenge is therefore to mitigate its impacts by reducing population abundance to protect biodiversity, while minimizing socio-economic damage to artisanal fisheries and aquaculture. Among the international proposed measures are intensive control fishing campaigns targeting key stages of the crab’s life cycle (UNEP/MAP-SPA/RAC 2025). The recommendations outlined in this report aim to support Mediterranean countries in adapting these measures to their own socio-economic and ecological contexts. They are organized around five strategic axes, both national and international, with the aim of structuring and strengthening collective and specific actions within a common Mediterranean control strategy to ensure greater effectiveness. The integration of acoustic telemetry data has refined the identification of priority intervention zones and periods. In alignment with international recommendations for the management of *C. sapidus* in the Mediterranean Sea (UNEP/MAP-SPA/RAC 2025), this study addresses the need to better understand the behaviour and ecology of this species in its invasive range. It also provides valuable new insights into how this invasive species utilizes Mediterranean lagoon ecosystems.

### 4.6. Conclusion and perspectives

This study provides the first detailed behavioural analysis of *Callinectes sapidus* in a Mediterranean lagoon, revealing both expected and novel patterns of activity, movement, and habitat use in its invasive range. The results highlight the species’ remarkable behavioural plasticity, particularly its ability to sustain long-distance movements and adapt to low tidal and variable salinity conditions. Notably, sex-based differences in behaviour and spatial distribution suggest reproductive and ecological strategies that may influence population dynamics and invasion success. These insights are critical for informing targeted management actions, such as the timing and location of control efforts. The integration of acoustic telemetry with ecological monitoring has proven effective in identifying key behavioural traits and priority intervention zones. This research directly contributed to Corsica’s first Territorial Control Plan for marine invasive species, offering a model for adaptive management in other Mediterranean regions. Future research should expand on these findings with longer tracking periods and broader sampling to better understand seasonal and ontogenetic shifts in behaviour.

## Acknowledgments

We sincerely thank fisherman Jean-Louis Guaitella and his team for their invaluable contribution to the blue crab sampling in the Biguglia Lagoon. We are also grateful to the administrative authorities for granting permission to conduct this study. Special thanks go to the Stella Mare platform for maintaining the receiver network at sea as part of the Corsic’Ange project, and to Caroline Bousquet for her coordination of the data collection. We extend our appreciation to Johann Mourier for his earlier guidance on the telemetry setup. We acknowledge the support of the UMR Sciences For Environment, particularly the Management of Mediterranean Waters research team led by Professor Vanina Pasqualini, for funding the Master’s grant awarded to K. Le Corre. We also thank Marina Luccioni and Dr. Nathalie Malet (Ifremer) for their assistance during fieldwork. Finally, we warmly thank the Stella Mare team, including technician Jean-Guy Vivoni for his help in constructing tag attachments, PhD student Marie-Catherine Raffalli, and Research Engineer Dr. Viviani Ligorini for their valuable support. We would like to thank the two anonymous reviewers for their constructive comments and suggestions, which helped improve the manuscript.

## Declarations

### Ethics approval

All applicable institutional and/or national guidelines for the care and use of animals were followed.

### Consent to participate

Not applicable

### Consent for publication

Not applicable

### Availability of data and material

Data have been deposited on github repository and are currently available as a private repository for the reviewing process on request: https://github.com/edhdurieux/Blue_crab_telemetry_Mediterranean_Corsica.git

### Code availability

Codes have been deposited on github repository and are currently available as a private repository for the reviewing process on request: https://github.com/edhdurieux/Blue_crab_telemetry_Mediterranean_Corsica.git

### Authors’ contributions

- **Eric D.H. Durieux**: Conceptualization (equal), Data curation (lead), Formal analysis (lead), Funding acquisition (equal), Investigation (equal), Methodology (equal), Project administration, Resources (equal), Software (lead), Supervision (equal), Validation, Visualization (equal), Original draft preparation (lead), Writing – Review & Editing (lead)
- **Klervi Le Corre**: Conceptualization (equal), Data curation (lead), Formal analysis (lead), Software (lead), Validation (equal), Visualization (equal), Original draft preparation (lead), Writing – Review & Editing (lead)
- **Guillaume Marchessaux**: Investigation, Original draft preparation, Writing – Review & Editing
- **Dimitri Veyssiere**: Investigation, Original draft preparation, Writing – Review & Editing
- **Sabrina Etourneau**: Investigation, Original draft preparation, Writing – Review & Editing
- **Marie Garrido**: Conceptualization (equal), Formal analysis, Funding acquisition (equal), Investigation (equal), Methodology (equal), Project administration (lead), Resources (equal), Supervision (equal), Validation, Visualization, Original draft preparation, Writing – Review & Editing

