## Supplementary materials for "Movements and habitat use of the invasive blue crab *Callinectes sapidus* in Mediterranean coastal lagoon"

**Supplementary material**

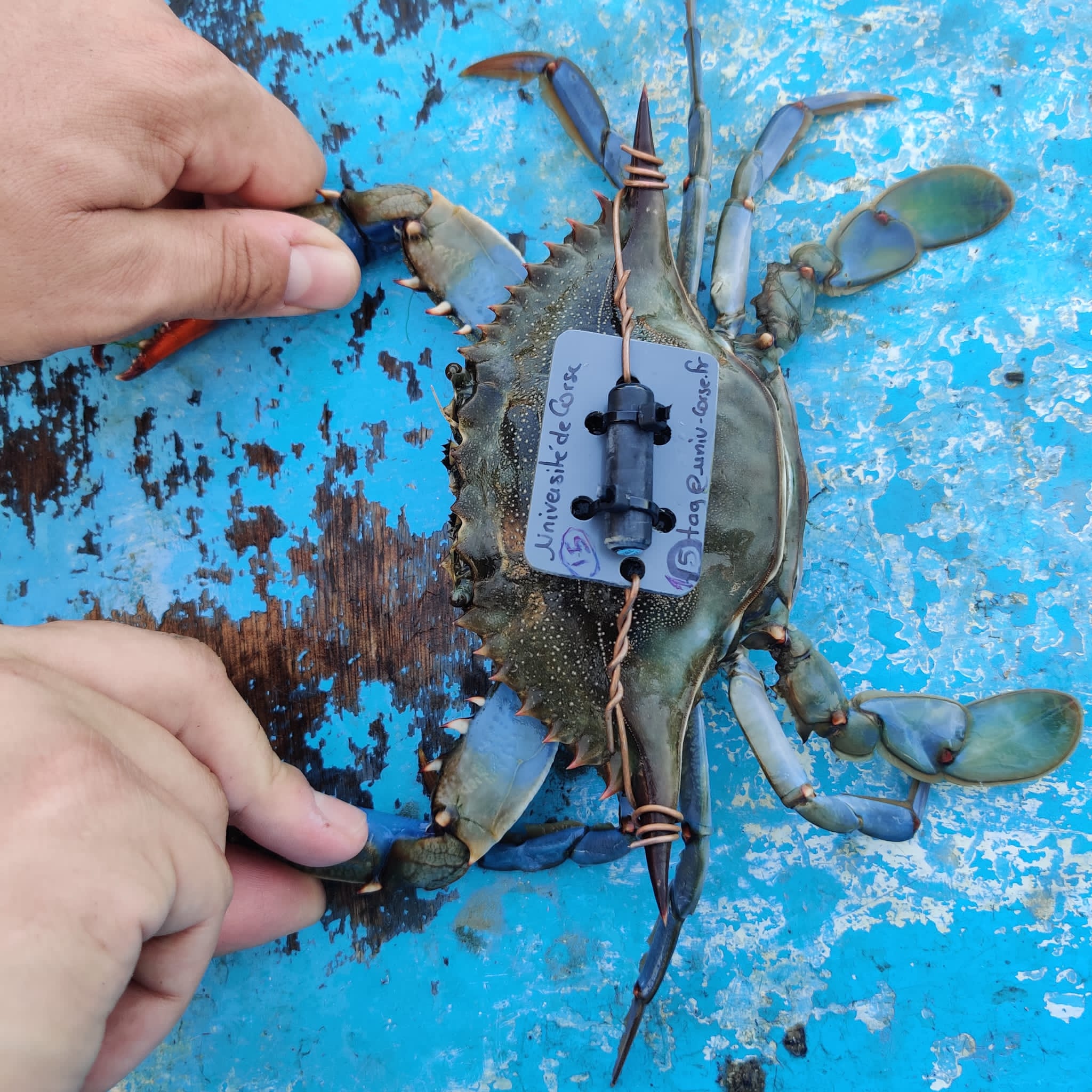

Supplementary Figure S1. Photograph of a female blue crab tagged with an acoustic transmitter for the telemetry study in the Biguglia lagoon (Corsica, France, Northwestern Mediterranean) in 2023. Photo by Marina Luccioni.

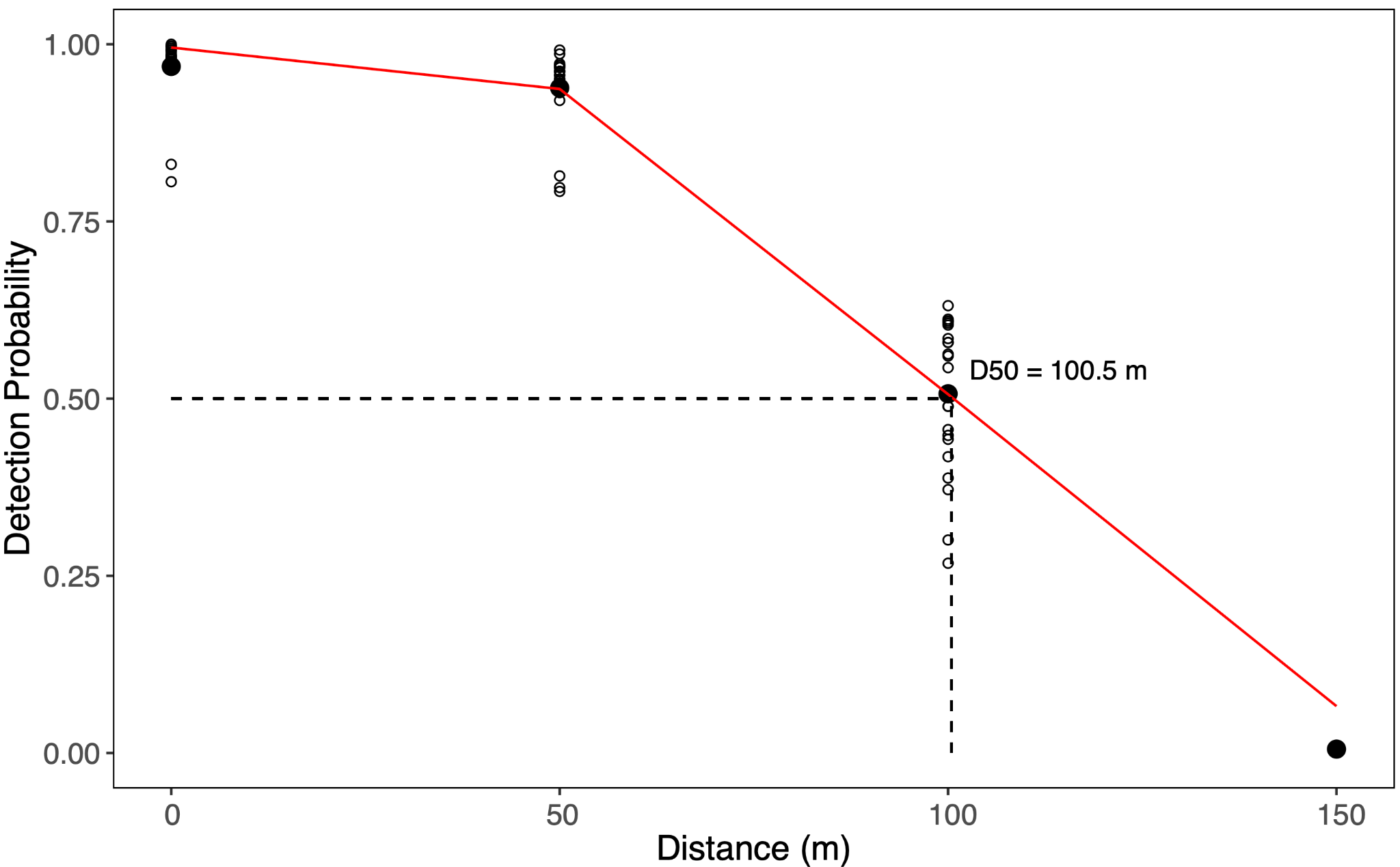

Supplementary Figure S2. Results of the range test carried out in the Biguglia lagoon in July 2024 using Thelma Biotel’s TBR700 receivers and ADT-MP9L tags. The white dots represent the average hourly detection rate, and the black dots represent the average detection rate for each transmitter distance (0, 50, 100 and 150 m).

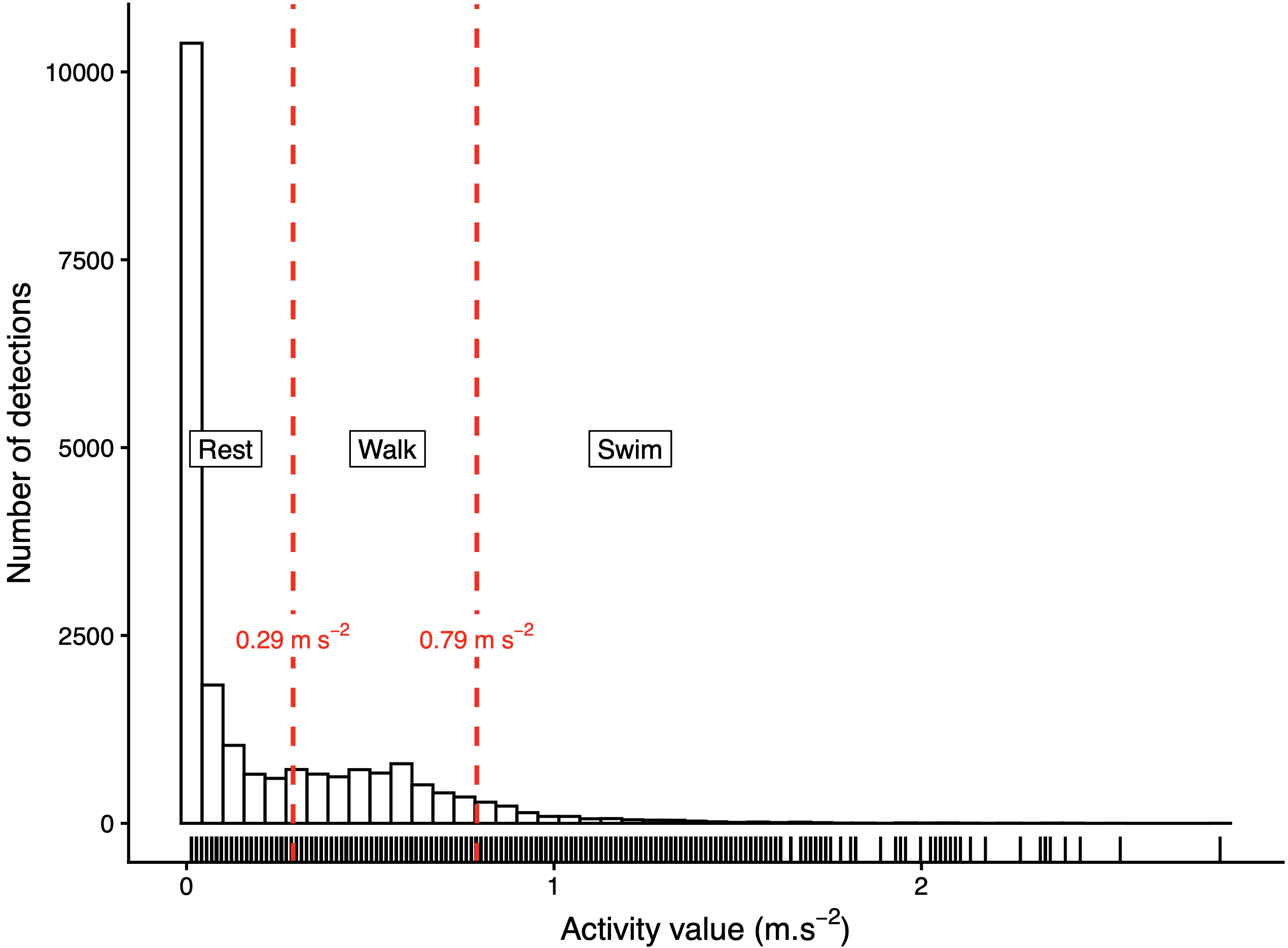

Supplementary Figure S3. Histogram showing the frequency distributions of activity of the blue crab (*Callinectes sapidus*) recorded during the acoustic telemetry survey in the Biguglia lagoon in 2023. Activity data were retrieved from acceleration transmitters (model ADT-MP9L, Thelma Biotel). Thresholds used to discriminate the three behaviors (rest, walk, swim) were defined following a K-means analysis.

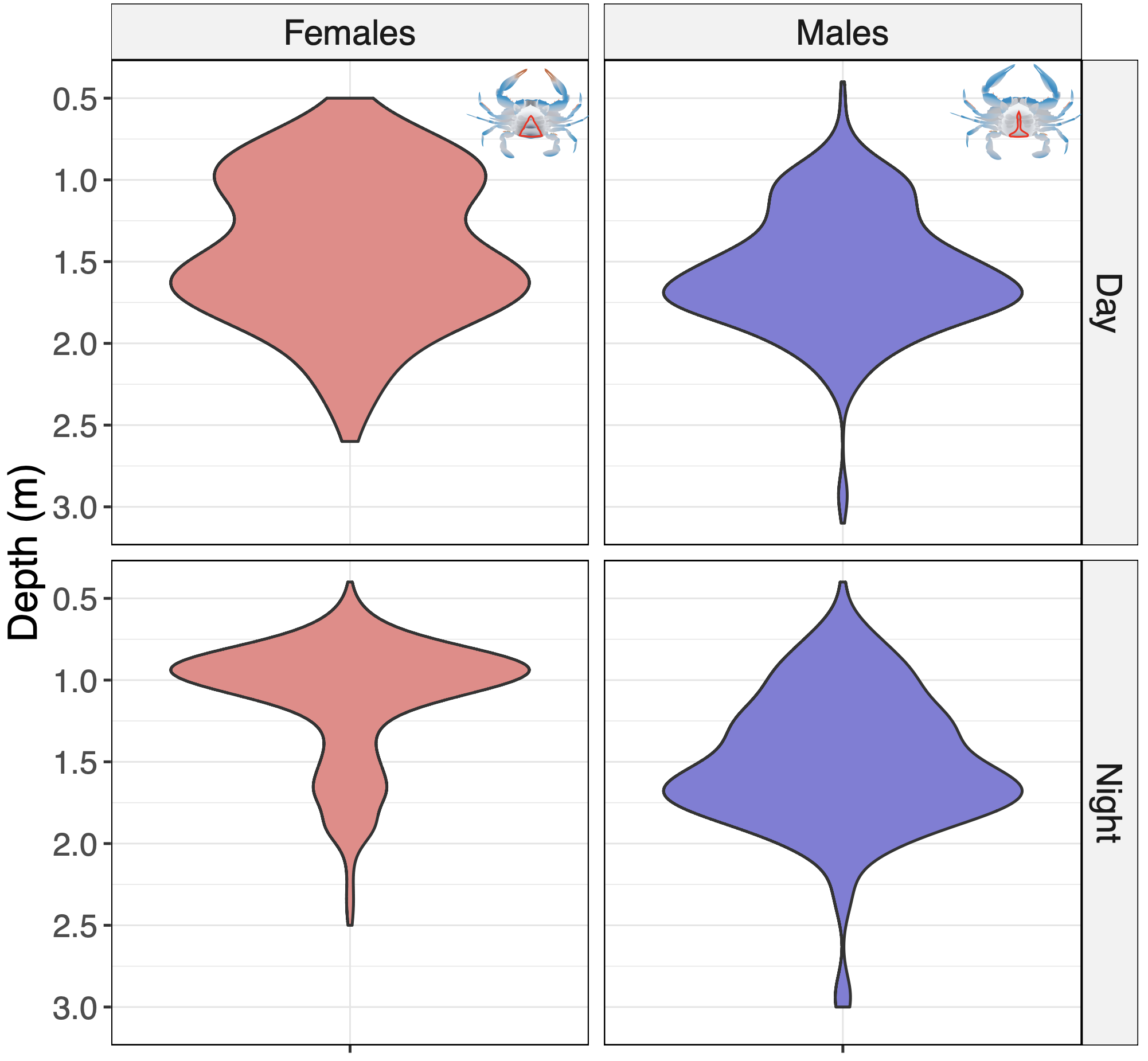

Supplementary Figure S4. Depth distribution by sex and daytime period of the blue crab (*Callinectes sapidus*) recorded during acoustic telemetry survey in the Biguglia lagoon in 2023.

Supplementary Table S1. Results of the generalized additive mixed model (GAMM) assessing the effects of sex, diel period (day/night), their interaction, carapace size, time, and individual variability on the blue crabs (*Callinectes sapidus*) recorded during acoustic telemetry survey in the Biguglia lagoon in 2023. Parametric coefficients, smooth-term statistics, and overall model fit metrics are presented. Abbreviations: SE, Standard Error; edf, Effective Degrees of Freedom; SD, standard deviation; AR(1), first-order autoregressive temporal correlation structure; φ, autoregressive parameter.

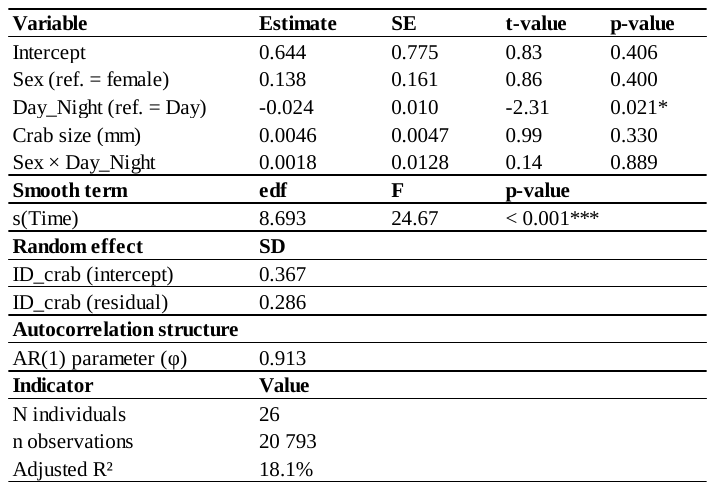
